# Phenotypic characterization of an Arabidopsis *stn7 stn8 pck2* triple mutant reveals only minor effects on chloroplast functions

**DOI:** 10.64898/2026.10.13.758734

**Authors:** Tim R. Demmig, Anja Rödiger, Kristin Peter, Johann Galonska, Birgit Agne, Omar Sandoval-Ibáñez, Sacha Baginsky

## Abstract

Reversible protein phosphorylation is a key regulator of chloroplast function. Although chloroplast kinases STN7, STN8, and plastid casein kinase 2 (pCK2) regulate photosynthetic acclimation and other chloroplast processes, their functional interplay remains poorly understood. We generated and characterized the first *Arabidopsis thaliana stn7 stn8 pck2* triple knockout mutant. Biochemical analyses confirmed the loss of all three kinase activities on established substrates. Growth resembled the *pck2* single mutant, whereas photosynthetic fluorescence parameters reflected the *stn7 stn8* double mutant. Thus, simultaneous disruption of all three kinases primarily combined the parental phenotypes, with little evidence of additive or synergistic effects under standard growth conditions. This triple mutant enables studies of chloroplast phosphorylation networks and demonstrates chloroplast physiology’s remarkable resilience to combined kinase loss.

## Introduction

Posttranslational modifications (PTMs) enable rapid modulation of protein activity in response to intra- and extracellular cues. Among these modifications, protein phosphorylation represents one of the best-characterized regulatory mechanisms and plays a pivotal role in rapid signal transduction, since it acts on already existing proteins **[1]**. In chloroplasts, the need for rapid regulation is particularly pronounced. Photosynthetic electron transport captures light energy to drive CO₂ fixation, yet light availability fluctuates constantly in natural environments. Efficient acclimation is therefore essential to prevent photodamage and maintain photosynthetic performance. Protein phosphorylation was recognized early as a key regulatory mechanism in this context with chlorophyll binding proteins ranging among the first candidates identified in the context of light dependent phosphorylation activity **[2]**.

Today it is known that the thylakoid-associated kinases STN7 and STN8 are central regulators of short-term light acclimation. STN7 primarily phosphorylates light-harvesting complex II (LHCII) proteins, thereby facilitating state transitions, a process that balances excitation energy distribution between photosystem II (PSII) and photosystem I (PSI) **[3]**,**[4]**,**[5]**. STN8 predominantly targets core proteins of PSII, including D1, D2, CP43 and PsbH and contributes to the dynamic turnover and repair of PSII, particularly during photoinhibitory stress **[6]**,**[7]**,**[8]**. Together, STN7- and STN8-mediated phosphorylation events regulate thylakoid membrane organization, photosystem function, and the maintenance of photosynthetic performance under variable environmental conditions **[4]**,**[7]**,**[9]**.

A functional connection between the thylakoid kinases STN7 and STN8 and members of the activity of bc1 complex kinase (ABC1K) family has recently been uncovered. Most chloroplast-localized ABC1K proteins are associated with plastoglobules, where they contribute to the regulation of prenyl-lipid metabolism, antioxidant homeostasis, and responses to environmental stress. ABC1K-mediated regulation of plastoquinone metabolism and chloroplast redox balance may influence the redox state of the photosynthetic electron transport chain, which is a key determinant of STN7 activation **[10]**. Consequently, ABC1Ks are thought to indirectly modulate thylakoid protein phosphorylation and photosynthetic acclimation **[11]**,**[12]**,**[13]**. Indeed, despite proposed functional roles, direct *in vivo* phosphorylation targets of these atypical kinases remain unidentified.

Beyond short-term acclimation, STN7 has also been implicated in long-term responses involving transcriptional regulation of photosynthesis-related genes, but no target proteins at the transcription machinery have been identified so far **[14]**,**[15]**,**[16]**,**[17]**. In contrast, direct phosphorylation of transcription-associated proteins has been demonstrated for plastid casein kinase 2 (pCK2), previously termed plastid transcription kinase (PTK) **[18]**,**[19]**,**[20]**. Recent studies revealed a broad spectrum of pCK2 substrates **[21]**,**[22]** comprising proteins involved in photosynthesis, RNA metabolism, protein folding and turnover, primary metabolism, and stress acclimation, indicating that pCK2 regulates multiple aspects of chloroplast function. The large number of pCK2-dependent phosphorylation events identified *in vivo* suggests that pCK2 contributes substantially to the organization and integration of chloroplast regulatory networks through reversible protein phosphorylation **[18]**,**[19]**,**[21]**,**[22]**.

While STN7 and STN8 primarily regulate photosynthetic acclimation through the phosphorylation of thylakoid proteins, pCK2 controls a much broader range of chloroplast processes via its diverse substrate spectrum. Despite increasing knowledge of their individual functions, little is known about the degree of functional overlap or crosstalk between these kinase systems. We therefore set out to characterize a *stn7 stn8 pck2* triple mutant (*3ko*) to investigate how thylakoid-associated signaling and broader plastid phosphorylation networks interact to maintain chloroplast function.

## Materials and Methods

### Plant lines and growth conditions

The Arabidopsis thaliana *stn7/stn8/pck2* triple knockout mutant (*3ko*) was generated by crossing the stn7/stn8 double mutant (SALK_073254/SALK_060869) with the *pck2* mutant (GK400A04), all in the Col-0 background. Homozygous plants were identified by PCR using gene- and T-DNA-specific primers (*stn7*: LB: 5’-GAGCTTGTGGGAATAGCTGTG-3’; RB: 5’-TAGTTGAACATGCGTGAGTCG-3’; *stn7* T-DNA-specific primers: Salk_LBb1.3: 5’-ATTTTGCCGATTTCGGAAC-3’; RB: 5’-TAGTTGAACATGCGTGAGTCG-3’; *stn8*: LB: 5’-TCTCCTTGAGCTTCTCTGCAG-3’; RB: 5’-GAATTCTTGGGGAGTAGGACG-3’; *stn8* T-DNA-specific primers: Salk_LBb1.3: 5’-ATTTTGCCGATTTCGGAAC-3’; RB: 5’-GAATTCTTGGGGAGTAGGACG-3’; *pck2*: LB:5’-GAATGAAGTGTGTTTCTGGCC-3’; RB: 5’-GGTAGGCGTTGAGTTCATCTG-3’; *pck2*T-DNA-specific primers: GABI_087474: 5’-ATAATAACGCTGCGGACATCTACATTTT-3’; FN50_GABI400A04: 5’-TTGACTTTTGAATCCACTAGCTT-3’). The *3ko* mutant, parental lines, and wild-type Col-0 controls were grown under short-day conditions (8 h light/16 h dark) at 190 µE, with day/night temperatures of 22/18°C. Unless used for PAM measurements, plants were cultivated on half-strength MS medium containing 0.8% sucrose and 1% agar. Plants for PAM measurements were grown on soil under the same light and temperature conditions at 65% relative humidity.

### Growth parameters

Shoot fresh weight and primary root length were assessed as general growth parameters. Shoot biomass was measured in 21-day-old plants using 30 individuals from three biological replicates. Plants were harvested directly into tubes to minimize water loss before weighing. Primary root length was determined in vertically grown, 14-day-old seedlings using ImageJ. Three biological replicates with 9–12 plants per genotype were analyzed. Statistical analysis was performed in GraphPad Prism using one-way ANOVA followed by Tukey’s multiple-comparison test, with p < 0.05 considered significant.

### Pigment determination

Chlorophyll *a*, chlorophyll *b*, carotenoids, and anthocyanins were quantified spectrophotometrically using seedlings from three biological replicates with five samples per replicate. For *pck2* and *3ko*, two seedlings were pooled per sample. Photosynthetic pigments were extracted from 21-day-old shoots with 80% acetone for 120 min at 4°C in the dark, according to Lichtenthaler and Buschmann **[23]**. Absorbance was measured at 470, 647, and 663 nm, and pigment contents were normalized to fresh weight. For anthocyanin analysis, frozen and homogenized plant material was extracted overnight in acidic methanol at 4°C. After chlorophyll removal with water and chloroform, the aqueous phase was measured at 530 and 657 nm. Anthocyanin content was normalized to fresh weight and expressed as A₅₃₀ − A₆₅₇ per gram fresh weight.

### Chlorophyll fluorescence measurements

6-week-old plants were dark-adapted for 30 min, and detached leaves were analyzed using a DUAL-PAM 100. Light-response curves were recorded by gradually increasing actinic light from 67 to 1,738 PAR, followed by a recovery phase at 33 PAR. Effective quantum yield [Y(II)], regulated non-photochemical energy dissipation [Y(NPQ)], and non-regulated energy dissipation [Y(NO)] were calculated according to Varotto et al. **[24]**. State transitions were analyzed using a modified protocol based on Pribil et al. **[25]**. After 30 min of dark adaptation, detached leaves were illuminated with red light for 15 min to induce state 2 and determine maximum fluorescence (F ₂). Leaves were then exposed to far-red light for 15 min to induce state 1. Saturating pulses were applied every 5 min to determine F ₁ and calculate qT

### Chloroplast isolation for protein analysis

Chloroplasts from 21-day-old *3ko* and wildtype seedlings were isolated in three biological replicates from enzymatically released protoplasts, following Fitzpatrick and Keegstra **[26]**. Seedlings were harvested at the end of the dark period, chopped, and washed in ice-cold isolation buffer containing 400 mM sorbitol, 20 mM MES-KOH (pH 5.2), and 0.5 mM CaCl₂. Cell walls were digested with 1.5% Cellulase Onozuka R-10 and 0.375% Macerozyme R-10 for 4 h at 22°C in the dark. All subsequent steps were performed at 4°C. Released protoplasts were filtered through a 200 µm nylon mesh, collected by centrifugation at 120 × g for 10 min, washed, and resuspended in chloroplast isolation buffer. Protoplasts were disrupted by passage through 20 and 10 µm nylon meshes. Intact chloroplasts were purified on a discontinuous 35%/85% Percoll gradient containing protease and phosphatase inhibitors by centrifugation at 2,600 × g for 20 min. Intact chloroplasts were collected from the interface above the 85% Percoll layer, pelleted at 700 × g, and washed in buffer containing 50 mM HEPES and 330 mM sorbitol. Protein concentrations were determined using the Bradford assay with RotiQuant and BSA as the standard. Chloroplast aliquots were stored at −80°C until proteomic analysis.

### SDS-PAGE & Immunoblotting

Plastid proteins (10 µg) were denatured in Laemmli buffer for 10 min at 75°C and separated by SDS-PAGE on 12% acrylamide gels according to Laemmli **[27]**. Gels were either stained with SYPRO Ruby as a loading control or transferred to PVDF membranes using a semi-dry three-buffer system. Proteins were detected by chemiluminescence using a Fusion FX7 imaging system. Primary antibodies against LHCB2, phosphorylated LHCB2, and RubisCO activase were obtained from Agrisera. The antibody against phosphorylated RubisCO activase at Thr78 was provided by Kim et al. **[21]**.

### Mass spectrometry

Chloroplast proteins (1 mg) were digested with trypsin after solubilization in NH₄HCO₃/RapiGest, reduction with DTT, and alkylation with iodoacetamide. Digestion was performed overnight at a trypsin-to-protein ratio of 1:20 and stopped with HCl. Of each sample, 100 µg was retained for total-proteome analysis and 900 µg was enriched for phosphopeptides using TiO₂ spin tips. Unenriched samples from *3ko* and wildtype plastids were analyzed in three biological and three technical replicates using nano-UPLC coupled to an Orbitrap Velos Pro. Phosphopeptide-enriched samples were analyzed in three biological replicates using an Orbitrap Elite. Peptides were separated on C18 columns with a 170 min acetonitrile gradient and measured by data-dependent acquisition of full MS scans followed by MS/MS of the 20 most intense precursor ions. Data were processed with MaxQuant 1.6.10.43/Andromeda against the *A. thaliana* Araport11 database. Protein-, peptide-spectrum-match-, and site-level FDRs were set to 1%. Carbamidomethylation was defined as a fixed modification; methionine oxidation, N-terminal acetylation, and serine/threonine phosphorylation were variable modifications. Protein abundance was quantified using MaxLFQ. Proteins detected in at least 5 of 9 measurements were analyzed after exclusion of contaminants, reverse sequences, and site-only identifications. Log₂-transformed intensities were compared using two-tailed t-tests with Benjamini–Hochberg correction. Significance required q < 0.05 and an absolute fold change ≥1.5. Phosphosites were considered reliable when detected in at least two of three biological replicates with a localization probability >0.75. Differential phosphorylation required p < 0.05 and an absolute fold change ≥1.5. Protein abundance data were used to distinguish phosphorylation changes from changes in total protein levels. SUBA5 **[28]**, MapMan4 **[29]**, and PhosPhAT 4.0 **[30]** were used for localization, pathway, and kinase-substrate analyses. WebLogo 3.9.1 was used for phosphopeptide motif analysis **[31]**.

## Results

### Phenotypic characterization

To investigate potential crosstalk between pCK2 and the thylakoid kinases STN7 and STN8, we crossed *pck2* knockout mutants with *stn7/stn8* double mutants to obtain triple knockout (*3ko*) lines lacking STN7, STN8, and pCK2 activity. Genetic and biochemical validation confirmed the absence of kinase activity, as evidenced by the loss of phosphorylation of established target proteins (e.g., LHCB2 and RCA) in the *3ko* lines (Supplemental Fig. 1 B). Phenotypic analyses were performed in a comparative manner with wildtype, *pck2* single and *stn7/stn8* double mutants. These analyses revealed no major defects in chloroplast biogenesis in any genotype (Fig. 1). However, growth was progressively reduced, with *pck2* and *3ko* plants exhibiting the most severe phenotype when compared with *stn7/stn8* double mutants (Fig. 1 A-C). Notably, *pck2* and *3ko* mutants displayed similarly impaired root growth as reported earlier for the single mutants **[22]** (Fig. 1 B and D). Taken together, the growth parameters reported here are largely dominated by the loss of *pck2* and our data do not present any evidence for a compounding effect on growth resulting from the loss of all three kinases compared to the *pck2* single mutant.

**Figure 1:**
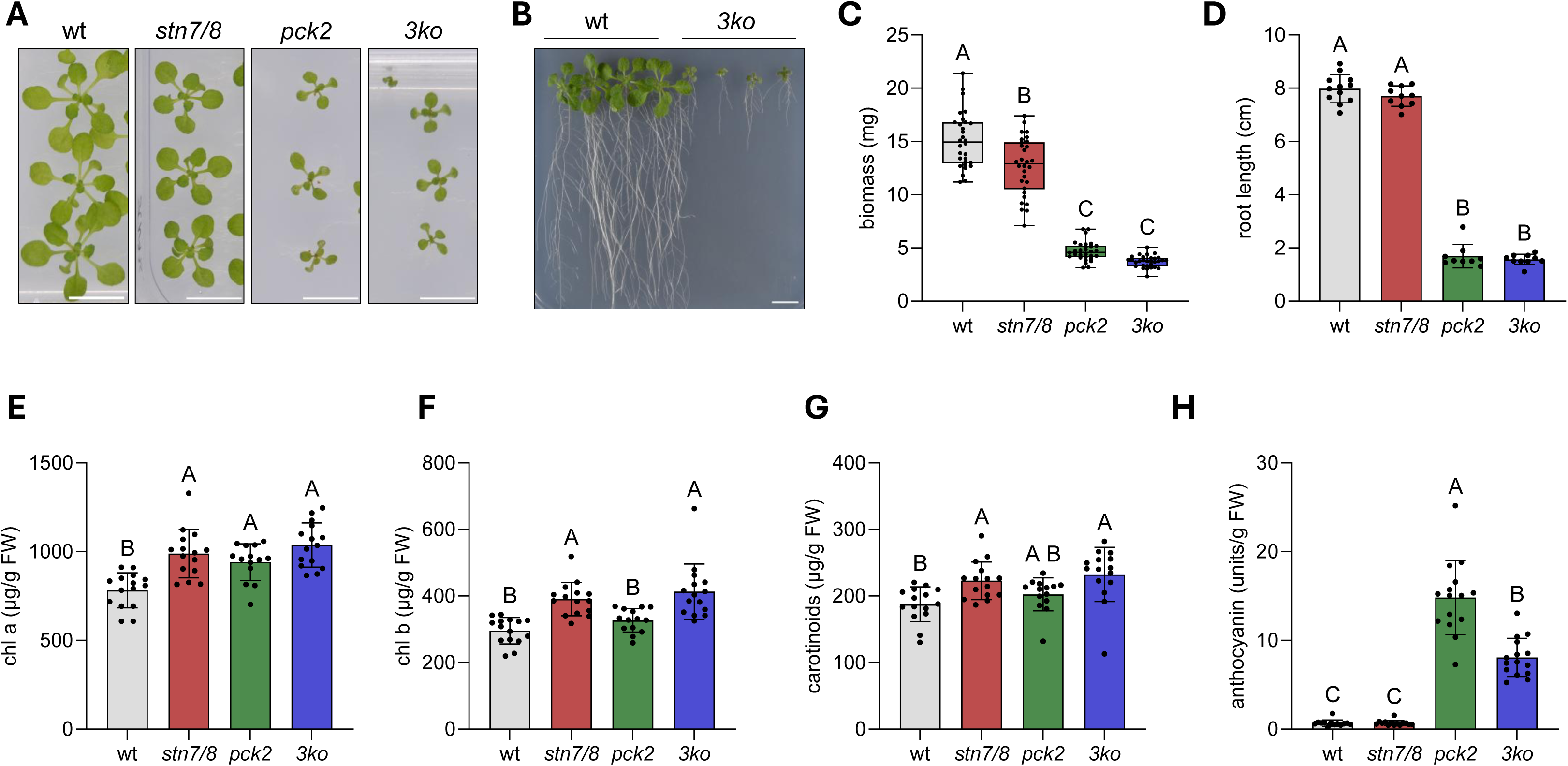
Characterization of protein kinase mutants. (A, B) Growth phenotypes of the *3ko* mutant, the parental lines used for crossing (*stn7/stn8* and *pck2*), and the wild type on ½ MS medium. Plants were grown for 21 days. Scale bar, 1 cm. (C) Shoot biomass determined after 21 days of growth. Box plots represent three biological replicates with 10 individuals each (*n* = 30). (D) Primary root length determined after 14 days using the freehand selection tool in ImageJ. Data are presented as the mean ± SD of three biological replicates (*n* = 9–12). (E–H) Spectrophotometric determination of pigment contents in 21-day-old plants. Pigment contents were normalized to fresh weight. Data are presented as the mean ± SD of three biological replicates with five individuals each (*n* = 15). Statistical significance was assessed using one-way ANOVA followed by Tukey’s multiple-comparison test. Differences were considered statistically significant at p-value < 0.05.

We next set out to measure pigment content to assess the effect of the triple knockout on the assembly of the photosynthetic machinery (Fig. 1 E-H).The Tukey post-hoc test revealed significant genotype-dependent differences in pigment content. Chlorophyll *a* was significantly lower in wildtype than in all three mutants that were indistinguishable from each other (Fig. 1 E). For chlorophyll *b* and carotenoids, wildtype and *pck2* showed that same trend, while the double and triple mutants accumulated significantly more of these pigments (Fig. 1 F and G). Thus, the photosynthetic pigments were stronger affected by the mutation in *stn7/stn8* compared to the *pck2* single mutant. Since the mutant lines lacking *pck2* appeared slightly darker green, we went ahead and measured anthocyanin levels. Anthocyanin levels were significantly higher in the *pck2* single and the *3ko* mutant suggesting some type of stress adaptation in the absence of pCK2 (Fig. 1 H).

Because anthocyanin biosynthesis is associated with stress responses, including response to photooxidative stress **[32]**, we assessed the photosynthetic performance by pulse amplitude modulation (PAM). Compared with the wildtype, *p*ck2 mutants exhibited only mild defects in effective quantum yield (Y(II)) under low-light conditions (Fig. 2A). However, the *3ko* plants displayed the most pronounced impairment, with the lowest values during the low-light illumination period and delayed recovery during the dark phase (Fig. 2A). These changes were more pronounced, yet comparable, to those observed in the *stn7/stn8* double mutant and are explained by changes in non-regulated energy dissipation (Y(NO)) and non-photochemical energy dissipation Y(NPQ) (Fig. 2 B, C). These results suggest mild PSII photodamage under low light transferred to the STN7/STN8 knock out, and indicate a possible functional interdependence among the three kinases. Therefore, and contrary to previous assumptions based on altered PsbH phosphorylation in *Chlamydomonas reinhardtii* **[33]**, the absence of STN7, STN8, and pCK2 did not cause a severe defect in photoinhibition recovery. Similarly, mild defects for the state transition kinetics were observed in *pck2* mutants compared to the wildtype, whereas similar kinetics were observed between *stn7/stn8* and *3ko* mutants (Fig. 2 D).Together, these findings suggest that the higher anthocyanin contents observed for *pck2* with alleviation in the 3ko mutants are unlikely to result solely from photooxidative stress, but may indicate underlying metabolic alterations **[34]**.

**Figure 2:**
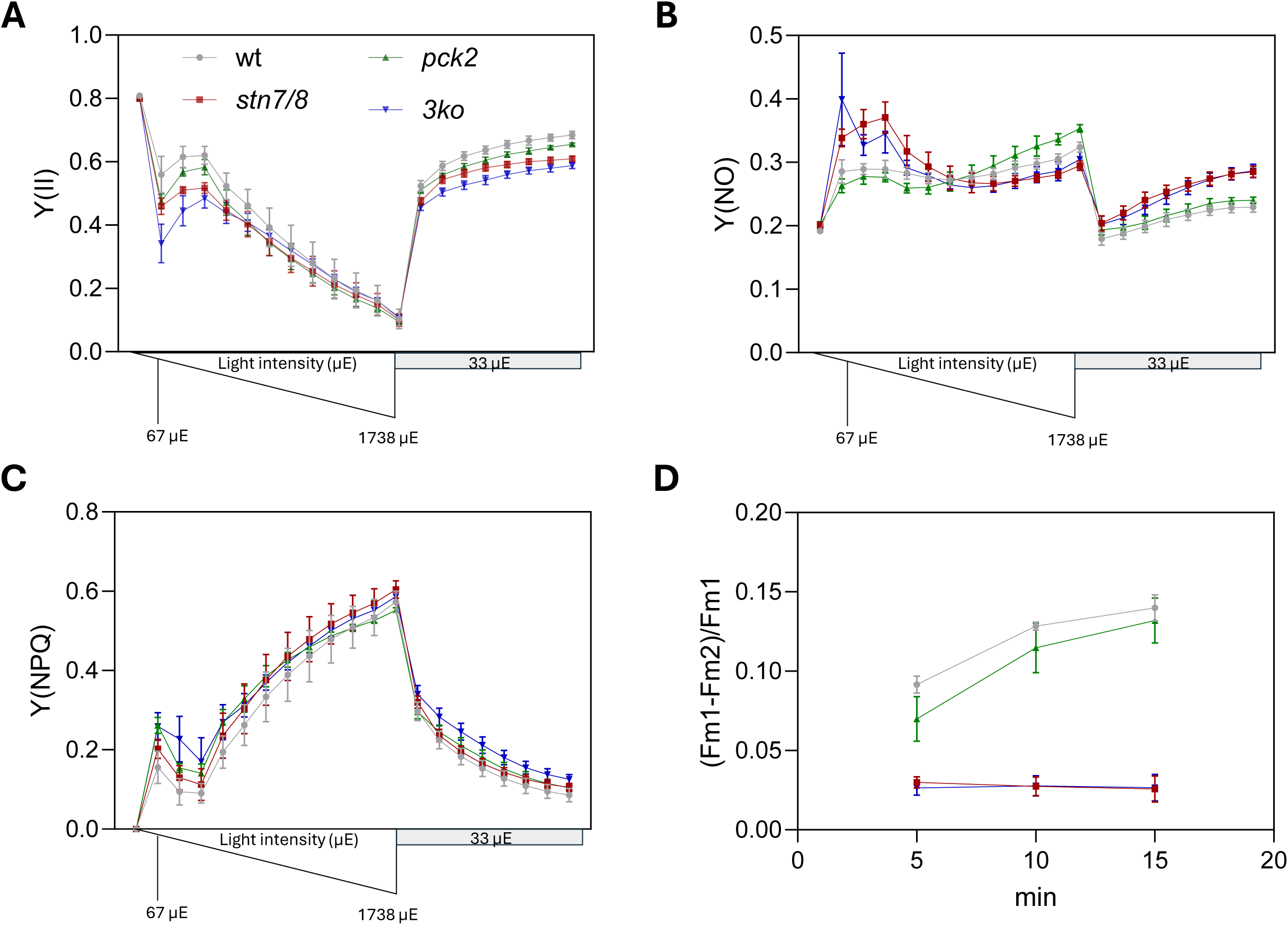
Analysis of photosynthetic performance using pulse-amplitude-modulation (PAM) chlorophyll fluorescence measurements. Plants were grown on soil for 42 days prior to measurement. (A-C) Photosystem II performance represented by the effective quantum yield of PSII (Y(II)), the quantum yield of non-regulated energy dissipation (Y(NO)) and the quantum yield of regulated energy dissipation (Y(NPQ)), respectively. Measurements were recorded over an 8-min course of increasing light intensity, ranging from 67 to 1738 µE, followed by a recovery phase at 33 µE. Each data point represents the mean ± SD of six plants measured at the respective light intensity. (D) Saturating-pulse analysis of the state transition from state 2 to state 1 over a period of 15 min. Data represents mean ± SD of six plants.

### Proteome and phosphoproteome analyses

To assess proteome adaptations to the triple kinase knockout, we performed quantitative proteome profiling on isolated chloroplasts from both genotypes in three biological replicates. Altogether, 1321 proteins were identified from wildtype and 1124 proteins from the *3ko* lines (Fig. 3 A). Applying strict criteria for significance including multiple testing corrections, only a relatively small subset of proteins is significantly changed (Fig. 3 C). Among 35 proteins classified as plastid, a significant upregulation of Calvin cycle enzymes such as phosphoglycerate kinase and GAP-DH regulatory protein is most pronounced, while RubisCO LSU is down to 82% of wild type (Fig. 3 F; Supplemental Fig. 2 and 4). Other proteins involved in electron transport such as some PSII subunits and NDH complex members are also upregulated, while enzymes involved in fatty acid synthesis are down regulated (Fig. 3 F; Supplemental Fig. 2). Interestingly, components of the light-harvesting machinery are rather down in the mutant, changing the PS II/LHCII ratio significantly to a high ratio in the *3ko* compared to wildtype (Fig. 3 E). This effect is similar, but weaker for photosystem I (Fig. 3 D). Overall, the proteomic data indicate a selective remodeling of the chloroplast proteome in the triple knockout mutant. The increased abundance of Calvin cycle enzymes and components of the photosynthetic electron transport chain may represent a compensatory response to altered carbon or energy metabolism. In contrast, the reduced abundance of LSU RubisCO and fatty acid synthesis enzymes suggests a redistribution of chloroplast resources away from carbon fixation and lipid biosynthesis toward maintaining photosynthesis and redox balance.

**Figure 3:**
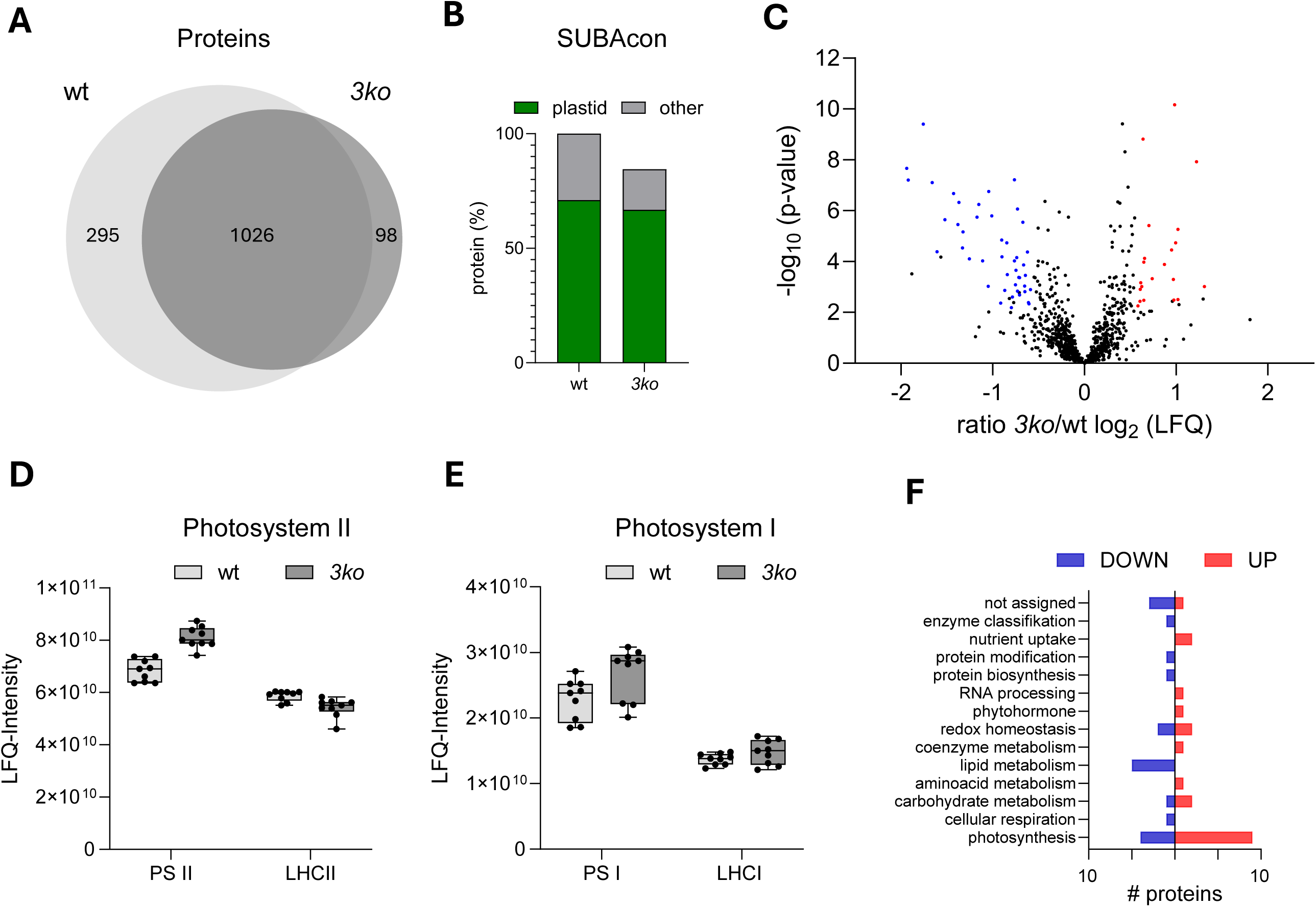
Comparative proteomic analysis of chloroplasts from the *3ko* mutant and wild type. The analysis was based on three biological replicates, each comprising three technical replicates. (A) Venn diagram showing the overlap between proteins detected in wild-type and *3ko* chloroplast samples. (B) Subcellular localization of proteins detected in the chloroplast samples, as predicted using the SUBA5 consensus classifier. (C) Volcano plot of all detected proteins, highlighting proteins with significantly altered abundance in the *3ko* mutant relative to the wild type. Significantly less and more abundant proteins are shown in blue and red, respectively. Differential abundance was defined by a Benjamini–Hochberg-adjusted p-value < 0.05 and a change in abundance of at least 1.5-fold. (D) Relative abundance of PSII and LHCII components in wild-type and *3ko* chloroplasts. Box plots show the summed LFQ intensities of all detected PSII and LHCII components from three biological replicates, each comprising three technical replicates. (E) Relative abundance of PSI and LHCI components in wild-type and *3ko* chloroplasts. (F) Pathway enrichment analysis of significantly altered plastid-localized proteins using the MapMan4 dataset. Proteins were included based on plastid localization according to the SUBA5 consensus classifier, a Benjamini–Hochberg-adjusted *P* value < 0.05, and a change in abundance of at least 1.5-fold.

**Figure 4:**
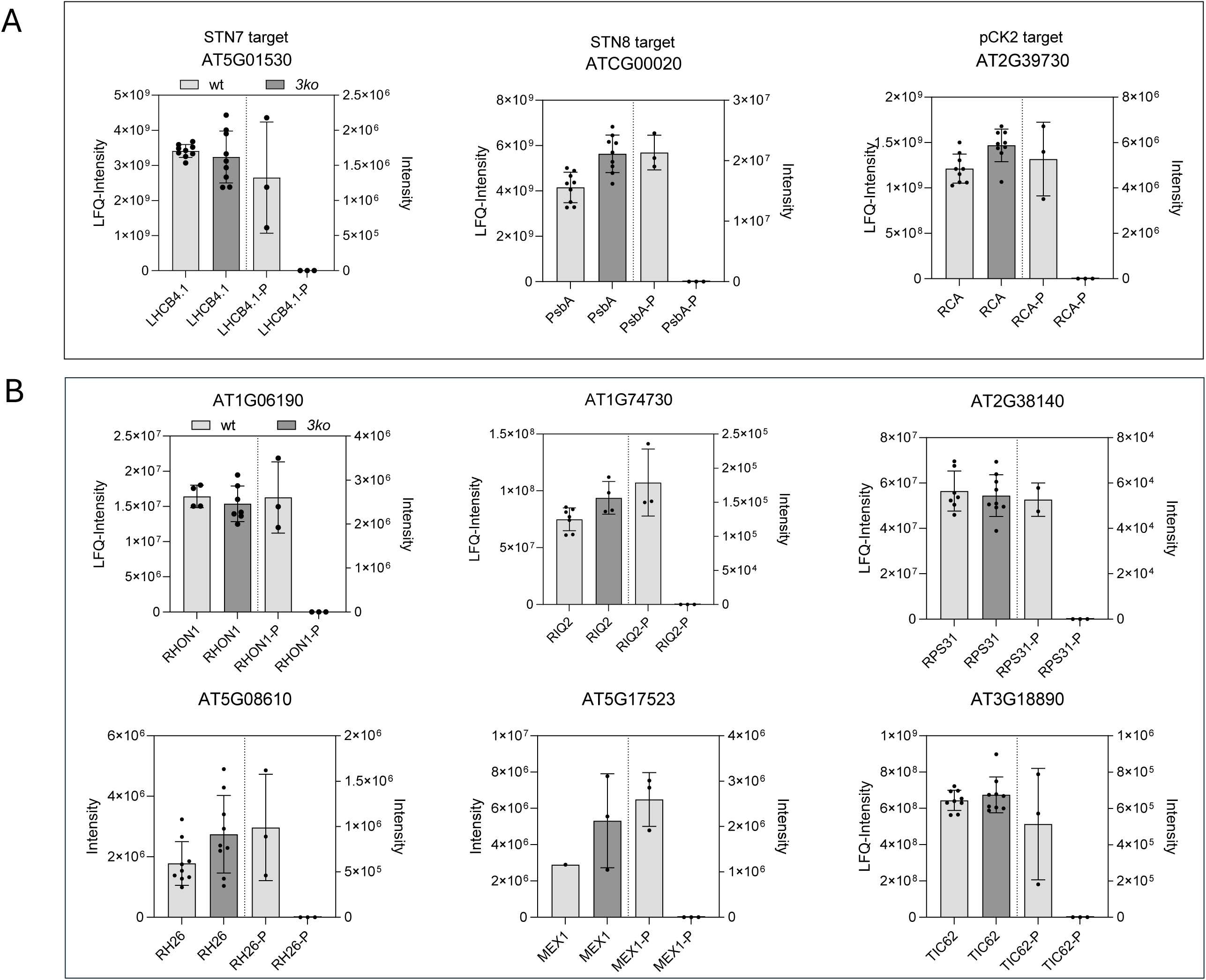
Comparative phosphoproteomic analysis of chloroplasts from *3ko* and wild-type plants. In each graph, the bars on the left show protein abundance (left *y*-axis), determined from chloroplast samples without prior phosphopeptide enrichment, whereas the bars on the right show phosphopeptide intensities (right *y*-axis), determined in phosphopeptide-enriched chloroplast samples. Protein abundances are presented as the mean ± SD of three biological replicates with three technical measurements each, and phosphopeptide intensities are presented as the mean ± SD of three biological replicates. Individual measurements are indicated by black dots. (A) Known targets of STN7, STN8, and pCK2, included as controls. (B) Phosphopeptides exhibiting patterns similar to those of the control targets but not previously reported as targets of STN7, STN8, or pCK2. Candidate phosphopeptides were required to be detected in at least two of the three wild-type biological replicates and to be undetectable in the *3ko* mutant.

The list of phosphorylation targets is in part an additive spectrum of substrates composed of known targets of STN7/STN8 and pCK2 **[35]**,**[22]** (Supplemental Fig. 3-5). However, several proteins have not been reported earlier as targets of the three different kinases (Fig. 4 B). Among them the nuclease RHON 1, the ribosomal proteins RPS31 and RPL12C, TIC62, EMB1865 and the plastid division protein PDV2. Notably, the maltose exporter 1 (MEX1) is among the newly identified targets, which is essential for the export of the starch degradation product maltose **[36]**. We also identified RIQ2 among our target list, which was recently described as a target of STN7 during leaf senescence **[37]**. Despite the loss of three important chloroplast kinases, several phosphorylation events in chloroplasts are still observable, some of which with higher abundance in the *3ko* line compared to wildtype, or even exclusively in *3ko*. This suggests that regulatory phosphorylations are catalyzed by additional chloroplast kinases. One interesting target protein here is fructose-bisphosphate aldolase 1 (At2g21330), that was identified in all replicates with *3ko* and never in wildtype. The phosphorylation site is in the peptide LA**<u>pS</u>**IGLENTEANR and does not conform with phosphorylation motifs of known protein kinases. This phosphorylation site is conserved, and its utilization was reported to be induced by drought and heat stress in maize leaves **[38]**.

We next aligned the phosphorylation sites of the identified target proteins that are not phosphorylated in the triple knockout mutants. Here the known and established pCK2 motif with the enrichment of acidic amino acids 1—3 upstream of the phosphorylation site is dominant (Fig. 5 A). The fact that threonine and serine are occurring at similar frequency in the aligned phosphorylation sites indicates activity of STN7 and STN8, with a slight overrepresentation of alanine and glycine directly downstream (-1), which is consistent with a motif that has been reported by Schönberg and colleagues for the two thylakoid kinases (Fig. 5 A) **[35]**. Thus, the alignment of phosphorylation sites underpins the observation that the target list is largely additive. In contrast, when those phosphorylation sites are aligned that are higher phosphorylated in the *3ko* plants compared to wildtype, an enrichment of proline in position +1 is observed, while glycine at position -1 and an acidic amino acid in position +3 are also observable (Fig. 5 B). Thus, our analysis reveals some of the characteristics of protein kinases other than STN7, STN8 and pCK2 acting in chloroplasts and shows that the chloroplast phosphorylation network can have different functional states, that apparently still result in proper chloroplast functioning.

**Figure 5:**
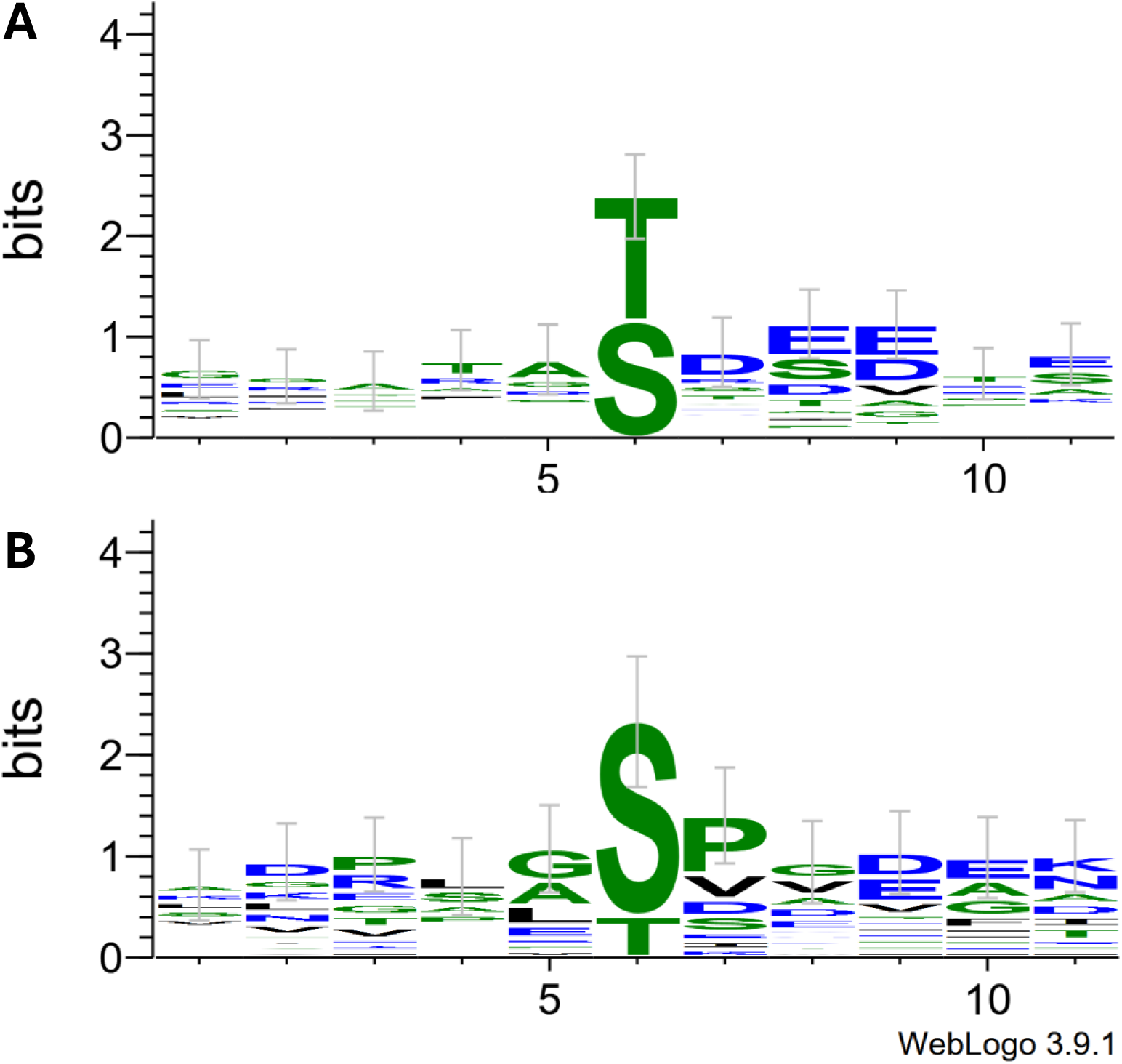
Sequence motifs surrounding plastid phosphorylation sites. Phosphorylation sites identified in this study were centered within sequence windows containing the five upstream and five downstream amino acid residues. Motif analysis was performed using WebLogo 3 with default settings. The height of each amino acid indicates its relative frequency at the corresponding position, and error bars represent Bayesian 95% confidence intervals. (A) Sequence motif surrounding phosphorylation sites not detected in the *3ko* mutant. (B) Sequence motif surrounding phosphorylation sites detected at a higher frequency in the *3ko* mutant compared to wild type.

## Discussion

Reversible protein phosphorylation is an important regulatory layer of chloroplast function, with STN7, STN8 and pCK2 representing three of the best-characterized plastid protein kinases in Arabidopsis **[35]**,**[22]**,**[39]**. Here, we combined loss of all three kinase activities in an *stn7 stn8 pck2* triple mutant to assess how simultaneous perturbation of these major phosphorylation pathways affects chloroplast function. Despite the loss of three major chloroplast kinases, the triple mutant was viable and developed functional chloroplasts. At the whole-plant level, the phenotype of the triple mutant was largely dominated by loss of pCK2. Reduced shoot growth and impaired root elongation were already reported for the *pck2* mutant and were not strongly aggravated by additional loss of STN7 and STN8 (Fig. 1) **[22]**. Similarly, alterations in state transitions characteristic for *stn7/stn8* plants were not further compounded by the loss of pCK2 activity (Fig. 2). Pigment accumulation was rather following the *stn7/stn8* genotype, so that a clear phenotypic distinction is visible between the different genotypes with growth parameters reflecting the *pck2* phenotype and photosynthetic fluorescence parameters and pigment accumulation reflecting the *stn7/stn8* double mutant phenotype. Thus, overall appearance of *3ko* plants reflected primarily a combination of parental phenotypes, while synergistic effects were not obvious at the whole plant level. A small additional effect of the combined kinase loss was observed for selected photosynthetic parameters (Fig. 2 A). Here, *3ko* performs worse compared to all other genotypes with lower PSII effective quantum yield at low light intensities and slower recovery kinetics from photoinhibition. However, this effect is rather small.

It is possible that long-term acclimatory remodeling of the chloroplast proteome partially buffers the loss of kinase-dependent regulation on photosynthesis. Photosynthetic parameters also do not explain the accumulation of anthocyanins in lines that lack pCK2 activity since anthocyanin accumulation is unlikely a result of photooxidative damage. Chloroplast carbon fixation, triose-phosphate export and cellular sugar availability have recently been shown to provide major signals for induction of anthocyanin biosynthesis during high-light acclimation **[40]**. Together with metabolic perturbations previously reported in *pck2* plants **[22]**, this raises the possibility that increased anthocyanin accumulation in *pck2* and *3ko* reflects changes in carbon and sugar metabolism rather than photosynthetic stress. These changes may be reflected in the proteome adaptations that we observed, particularly for Calvin cycle enzymes (Fig. 3; Supplemental Fig. 2). Furthermore, phosphorylation characteristics of fructose-bisphosphate aldolase 1 and MEX1 together with the observed proteome adaptations support adjustments in the wiring of carbohydrate metabolism (Fig 4; Supplemental Fig. 5). Whether or not this causes anthocyanin accumulation will require further research that uses e.g. large-scale metabolite profiling.

The phosphoproteomic data show that the molecular consequences of the triple knockout are more pronounced than its physiological phenotype. Phosphorylation sites lost in the *3ko* largely reflected the combined substrate specificities of STN7/STN8 and pCK2 (Fig. 4; Supplemental Fig. 3-5). In particular, the enrichment of acidic residues surrounding sites absent in the *3ko* mutant is consistent with the established pCK2 phosphorylation motif, whereas the occurrence of small residues such as glycine and alanine adjacent to other sites resembles sequence preferences previously reported for STN7/STN8-dependent phosphorylation **[35]**,**[22]**. These sequence motifs are compatible with the combined loss of established pCK2- and STN7/STN8-dependent phosphorylation events (Fig. 5 A). Importantly, the *3ko* mutant did not lose phosphorylation activity completely. A distinct set of sites showed increased phosphorylation, with a pronounced enrichment of proline at the +1 position and additional preferences for glycine and acidic residues (Fig. 5 B). This motif differs from those associated with STN7, STN8, and pCK2 and therefore suggests that other kinase activities contribute to the altered phosphorylation state of proteins in the *3ko* lines. It remains unclear which kinase(s) are active in the *3ko* mutant, and how this activity is integrated with regulatory dephosphorylations by phosphatases **[41]**,**[8]**.

Together, our data show that removal of three important kinases from chloroplasts induces a reorganization but not a general breakdown of the chloroplast phosphorylation network. The phosphorylation activity of other kinases may buffer the loss of phosphorylations by the three kinases investigated here and thus contribute to the relatively mild physiological phenotype of the triple mutant. Selective remodeling of photosynthetic protein abundance, changes in carbohydrate metabolism, and phosphorylation events mediated by remaining kinases or altered phosphatase activities may all contribute to this buffering. The *3ko* thus represents a valuable genetic resource for identifying residual components of the chloroplast phosphorylation network and for determining the limits of physiological compensation under more challenging environmental conditions.

## Acknowledgement

The authors are grateful for financial support from DFG grant BA 1902/2-4 and would like to thank Regina Oworah-Nkruma for technical support. Additionally, we thank Thomas Pfannschmidt for providing preliminary PAM data of the triple knockout mutant.

## Data availability

The mass spectrometry proteomics data have been deposited to the ProteomeXchange Consortium via the PRIDE (https://www.ebi.ac.uk/pride) partner repository with the dataset identifier PXD083272.

