## Supplemental Figures for "Phenotypic characterization of an Arabidopsis *stn7 stn8 pck2* triple mutant reveals only minor effects on chloroplast functions"

#### Suppl Figure 1

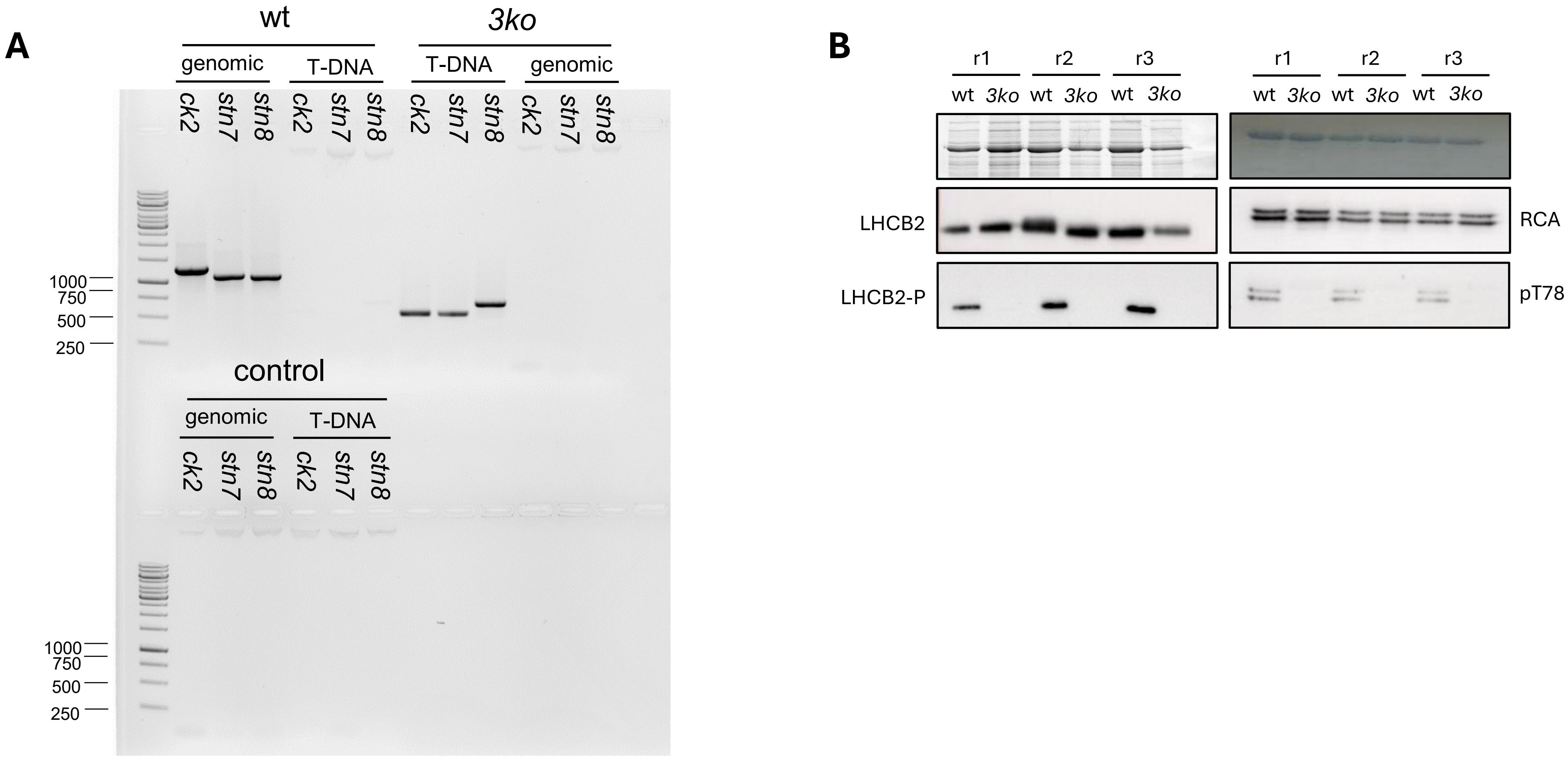


**Supplementary Figure 1**: Genetic and biochemical validation of the triple knockout mutant (*3ko*). (A) Representative PCR analysis of a homozygous triple knockout mutant and the corresponding wildtype control. Gene- and T-DNA-specific primer were used to confirm the knockout of STN7, STN8, and pCK2. (B) Western blot analysis of target protein phosphorylation. For STN7, LHCB2 and the corresponding phospho-specific antibody (Agrisera) were used. Rubisco activase (RCA; Agrisera) and a phospho-specific antibody recognizing threonine 78 (pT78; Kim et al. (2016)) were used to assess RCA phosphorylation as a control for pCK2 activity.

**Supplementary Figure 2:** Quantitative proteome analysis of isolated chloroplasts from the *3ko* mutant and wildtype plants. Changes in protein abundance are expressed as $log_2$ fold changes, with down- and upregulated proteins shown in blue and red, respectively. MapMan4 was used for pathway classification. The data comprises three biological replicates, each measured in three technical replicates. Only proteins classified as plastid-localized by SUBA5, passing the Benjamini–Hochberg correction ($q < 0.05$) and exhibiting a minimum absolute fold change of 1.5 were included.

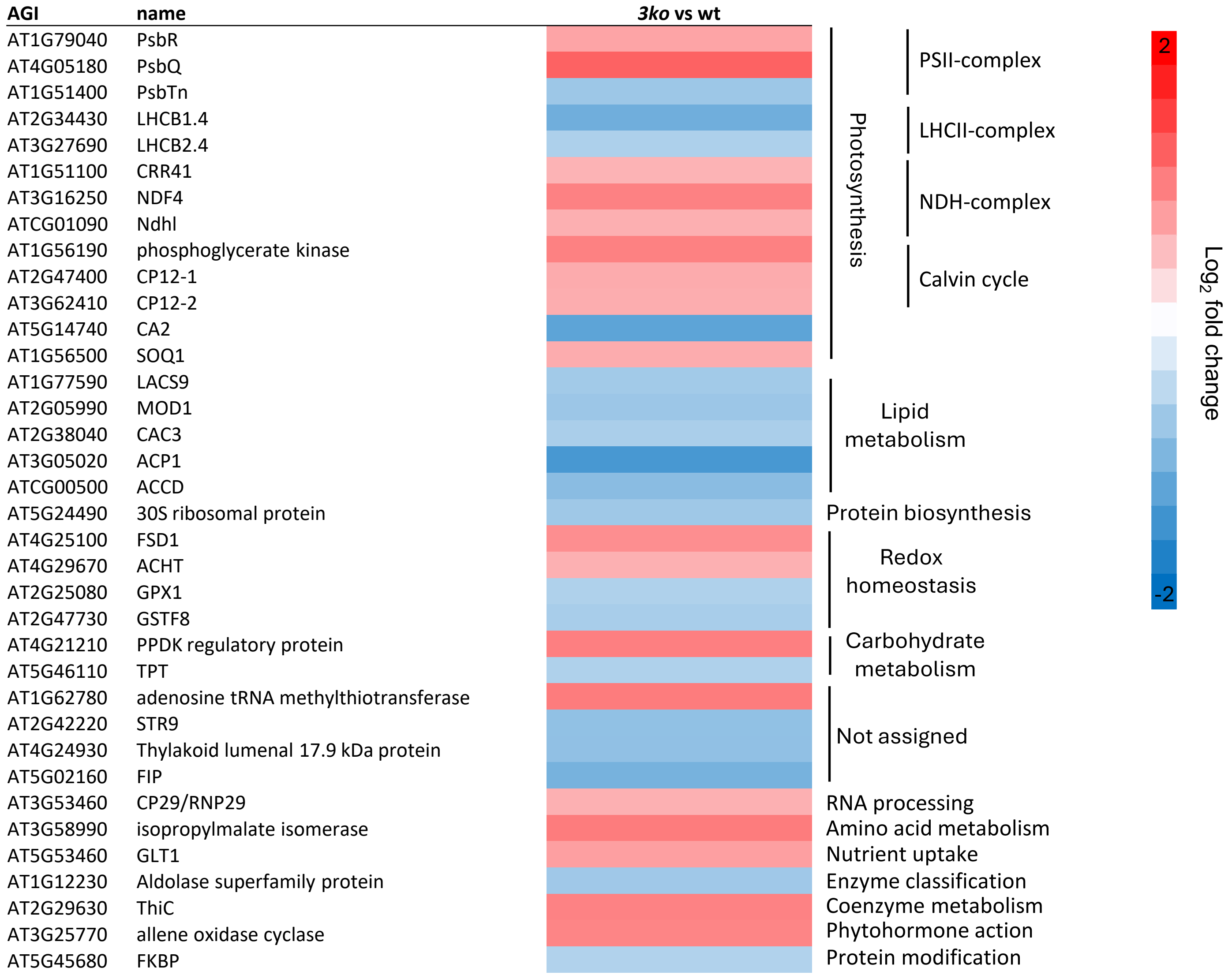

### Suppl Figure 3

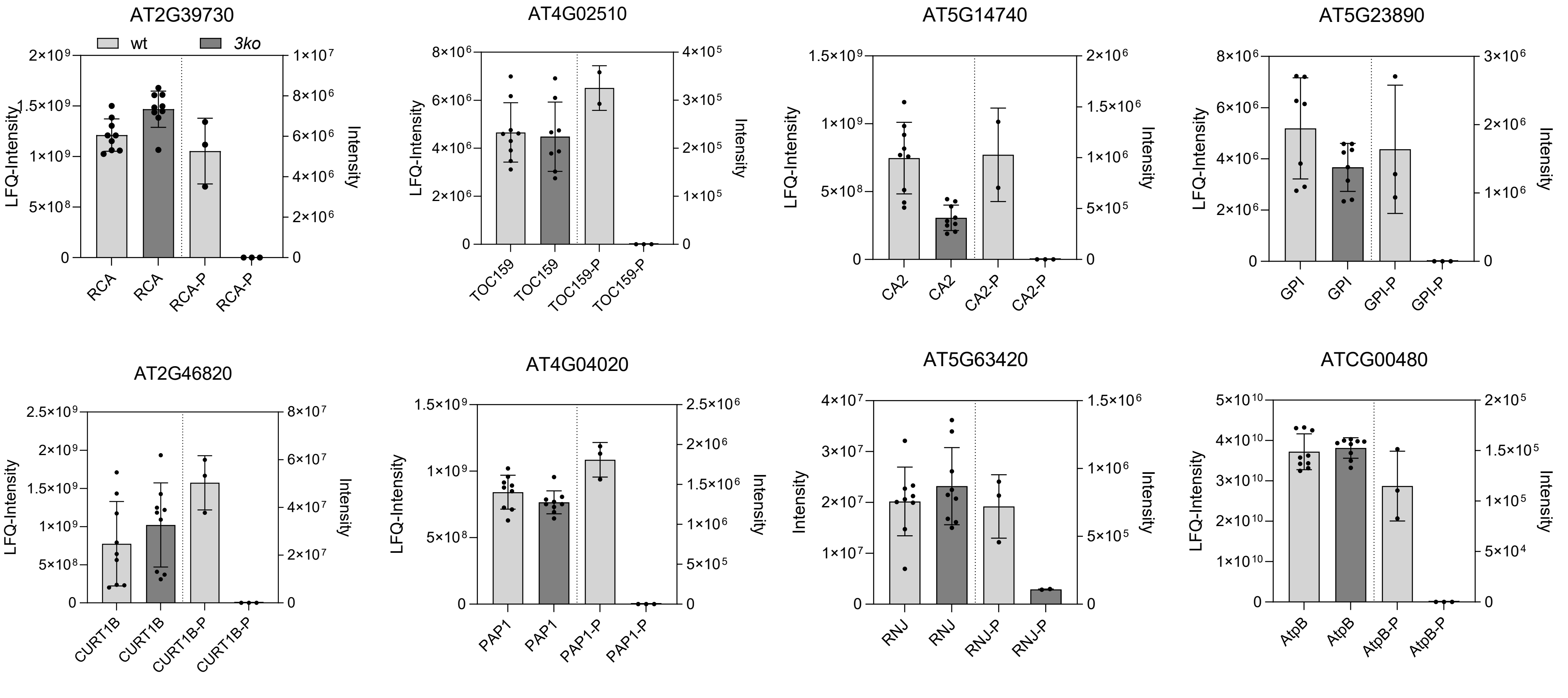


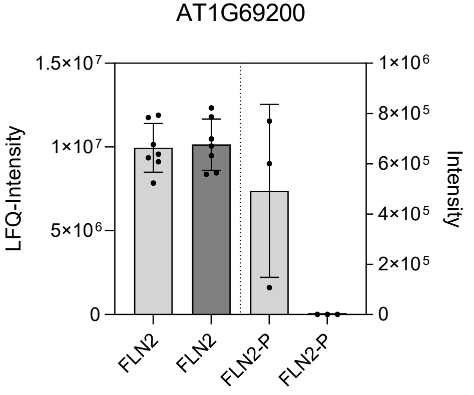


**Supplementary Figure 3**: Missing pCK2 target phosphorylation in the *3ko* mutant. Protein abundance (left y-axis) and phosphopeptide intensity (right y-axis) of pCK2 targets in wildtype and *3ko* plastids. Protein abundance was determined from unenriched plastid samples based on three biological replicates with three measurements each, whereas phosphopeptide intensities were determined from $TiO_2$-enriched plastid samples based on three biological replicates. Individual measurements are shown as black dots, columns represent mean values and error bars indicate standard deviation.

**Supplementary Figure 4:** Missing STN7/ STN8 target phosphorylation in the 3ko mutant. Protein abundance (left y-axis) and phosphopeptide intensity (right y-axis) are represented by mean ± SD and black dots represent individual measurments.

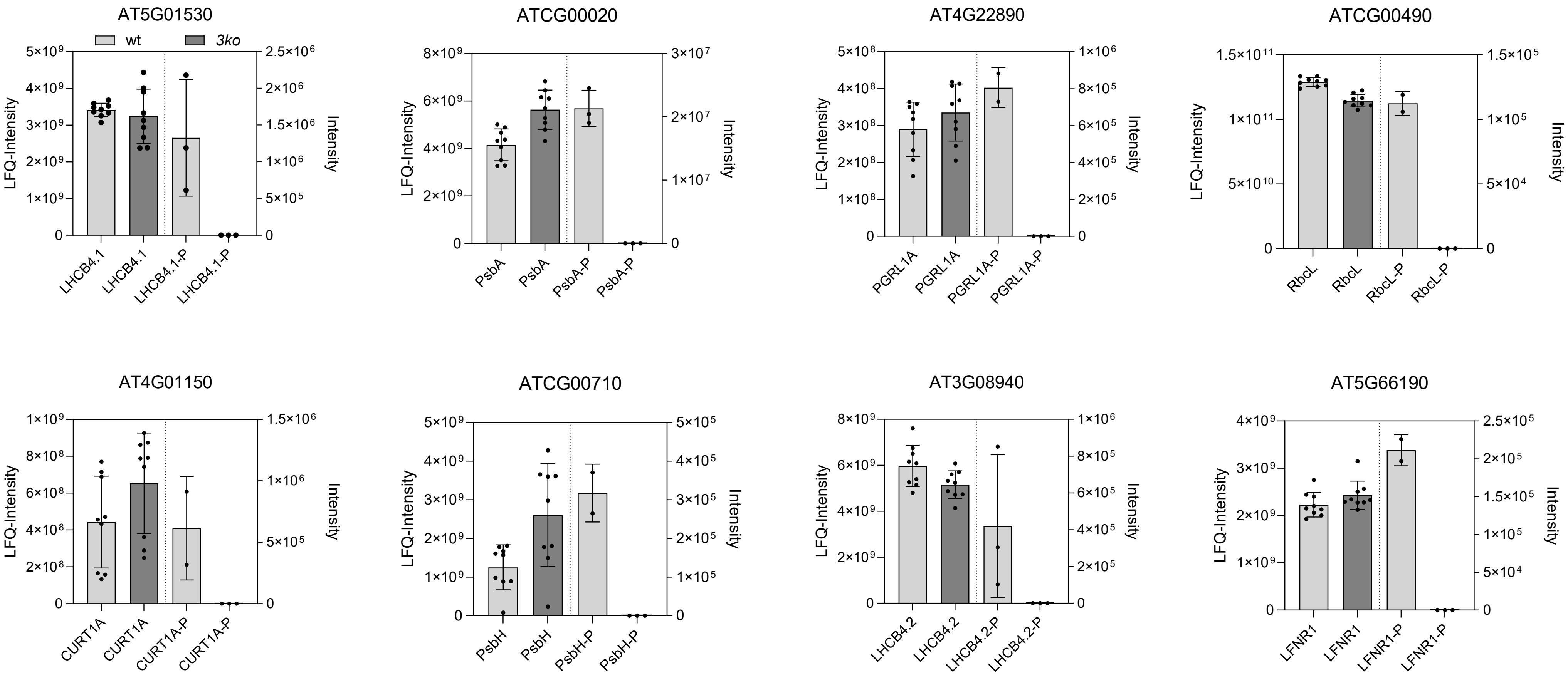

**Supplementary Figure 5:** Protein abundance (left y-axis) and phosphopeptide intensity (right y-axis) of selected proteins showing phosphorylation events exclusively detectable in the 3ko mutant compared to wildtype.

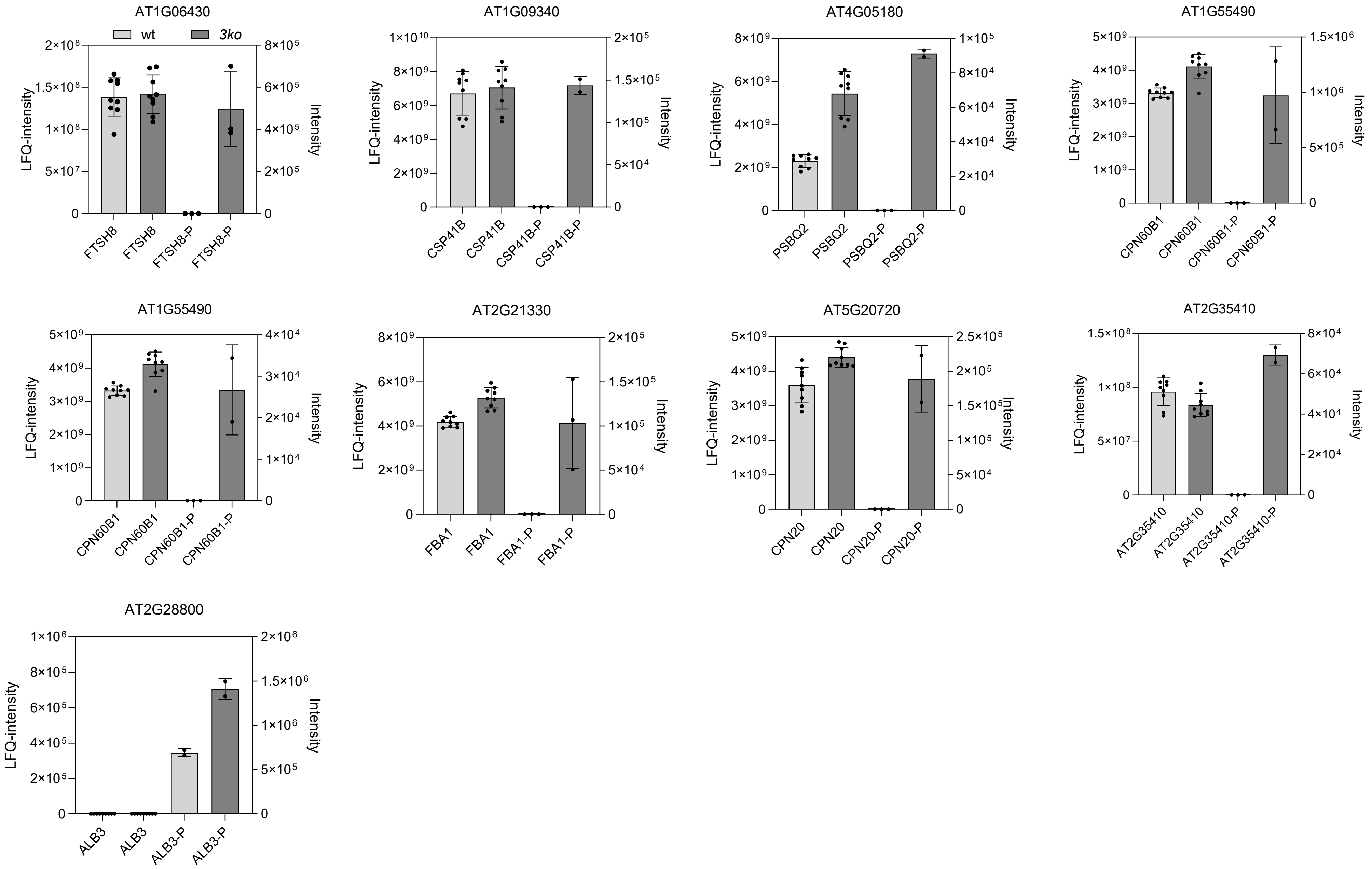
